# HIV Peptidome-Wide Association Study Reveals Patient-Specific Epitope Repertoires Associated with HIV Control

**DOI:** 10.1101/307314

**Authors:** Jatin Arora, Paul J. McLaren, Nimisha Chaturvedi, Mary Carrington, Jacques Fellay, Tobias L. Lenz

## Abstract

Genetic variation in the peptide-binding groove of the highly polymorphic human leukocyte antigen (HLA) class I molecules has repeatedly been associated with HIV-1 control and progression to AIDS, accounting for up to 12% of the variation in HIV-1 set point viral load (spVL). This suggests a key role in disease control for HLA presentation of HIV-1 epitopes to cytotoxic T cells. However, a comprehensive understanding of the relevant HLA-bound HIV epitopes is still elusive. Here we developed a peptidome-wide association study (PepWAS) approach that integrates HLA genotypes and spVL data from 6,311 HIV-infected patients to interrogate the entire HIV-1 proteome (3,252 unique peptides) for disease-relevant peptides. This PepWAS approach revealed a core set of epitopes associated with spVL, including previously characterized epitopes but also several novel disease-relevant peptides. More importantly, each patient presents only 16 (±7) and 6 (±6) of these core epitopes through their individual HLA-B and HLA-A variants, respectively. Differences in these patient-specific epitope repertoires account for almost all the variation in spVL previously associated with HLA genetic variation. PepWAS thus enables a comprehensive functional interpretation of the robust but little understood association between HLA and HIV-1 control, prioritizing a short and targetable list of disease-associated epitopes for personalized immunotherapy.

---

HLA class I proteins are thought to play a critical role in immune recognition of HIV-1 by presenting endogenously processed viral peptides at the surface of infected cells to cytotoxic T cells, in order to trigger destruction of the infected cells ^1^. Indeed, genetic variation in the HLA region has repeatedly been identified as the major genetic determinant of HIV-1 control in genome-wide association studies ^2,3^. Most recently, McLaren *et al.* ^4^ fine-mapped the entire HLA’s association with HIV-1 control and disease progression to five independent amino acid residues in the peptide binding groove of the HLA-B and HLA-A molecules. These five residues alone accounted for 12.3% of the variation in viral load, suggesting a major role for specific HLA-presented viral epitopes in HIV-1 control. However, our understanding of the disease-relevant viral epitopes is still incomplete, hampered by the unfeasible challenge of employing a full-factorial experimental assay to screen the entirety of the HIV-1 peptidome for binding by all relevant HLA alleles. Therefore, we developed a novel computational approach that identifies and prioritizes disease-associated peptides based on individual HLA genotype and disease phenotype information. Using a unique dataset of 6,311 individuals of European ancestry with chronic HIV-1 infection (**Table S1**) we screen the entire HIV-1 peptidome for epitopes that explain the known association between HLA genetic variation and HIV-1 control. This dataset from McLaren *et al.* ^4^ comprises pre-treatment level of set point viral load (spVL) as a correlate of disease progression ^5^ and imputed HLA genotypes (4-digit allele resolution). We focused on the two HLA loci (HLA-B and HLA-A) reported to have independent associations with HIV-1 control and disease progression ^4^. Potential HLA-bound peptides were identified using an established computational algorithm that is based on empirical training data ^6^ and integrates several complementary prediction methods in a consensus approach, outperforming comparable algorithms ^6,7^. Such algorithms have been used in a wide spectrum of HLA-related studies ranging from vaccine design to cancer evolution and HIV disease genetics ^8–10^. Without *a-priori* selection, we screened all possible 9mer HIV-1 peptides (N = 3,252) in a sliding window across the entire HIV-1 M group subtype B reference proteome ^11^ against all represented HLA-B and HLA-A alleles (344,712 HLA:peptide complexes), and identified 214 and 173 distinct HIV-1 peptides predicted to be bound by one or more of the represented HLA-B and HLA-A alleles, respectively.

In order to evaluate the significance of the predicted epitope repertoires, we interrogated several layers of empirical evidence. Firstly, the identified subsets of predicted HLA-B- and HLA-A-bound HIV-1 peptides were strongly enriched for experimentally tested cytotoxic T-lymphocyte (CTL) epitopes from the Los Alamos HIV Molecular Immunology Database ^12^ (HLA-B: OR = 5.8, *P* < 0.0001; HLA-A: OR = 5.0, *P* < 0.0001; **Figure 1A; Table S1 and S2**). Several predicted HLA-peptide complexes, e.g. B*57:01-IW9_15-23_, B*58:01-SW9_241-249_, in *Gag*, B*14:01-EL9_584-592_ in *Env* and B*57:01-HW9_116-124_, B*35:01-YY9_135-143_ in *Nef* were previously shown to be either frequently recognized by CD8^+^ T-cells ^13^ or associated with low viremia and slow disease progression ^14,15^. Secondly, we observed a negative correlation between an allele’s effect on spVL and the number of HIV-1 peptides that are predicted to be bound by that allele (Kendall’s tau = − 0. 26, *P* = 0.002; **Figure 1B; Table S3**). This observation supports the assumption that a greater number of HLA-presented peptides, together with the availability of a corresponding diverse TCR repertoire, confer greater protection against HIV ^10^. More importantly, this correlation indicates that our computationally predicted peptides are enriched for epitopes that contribute to the well-established association between HLA and viral load. No such association was observed when correlating the HIV-specific allele effect with binding to peptides from other viruses, confirming an HIV-1 peptide-specific effect (**Figure S2**). Thirdly, we used our prediction pipeline to identify HLA allele-specific *escape variants* in autologous HIV-1 sequences (available for a subset of patients, **Table 1**). Escape variants exhibit sequence variation that impedes either HLA binding or TCR binding to a given epitope and thus allow the virus to avoid CTL recognition. Here we focused on variants escaping HLA binding, expected to evolve in response to patient-specific HLA restriction ^16–18^. Focusing on the allele HLA-B*57:01, associated with strongest protection against HIV-1 ^16,19,20^, we tested whether autologous sequences of patients carrying B*57:01 harbored more escape variants specific to this allele than patients who did not carry this allele. Our autologous data covered 18 B*57:01-bound peptides, 12 of which showed no significant difference in the proportion of escape variants between B*57:01 carriers and non-carriers. Of the six HIV-1 peptides that did show a significant difference, all showed a higher proportion of escape variants in B*57:01 carriers (Chi-squared test, all *P* < 0.01 after Bonferroni correction; **Figure 1C**). This observation confirms significant B*57:01-specific restriction on HIV-1 and suggests that our prediction pipeline identifies HLA-relevant variation in HIV-1 peptides. Following these independent layers of evidence, we subsequently refer to these predicted HLA-bound peptides as *predicted epitopes*.

**Figure 1.**
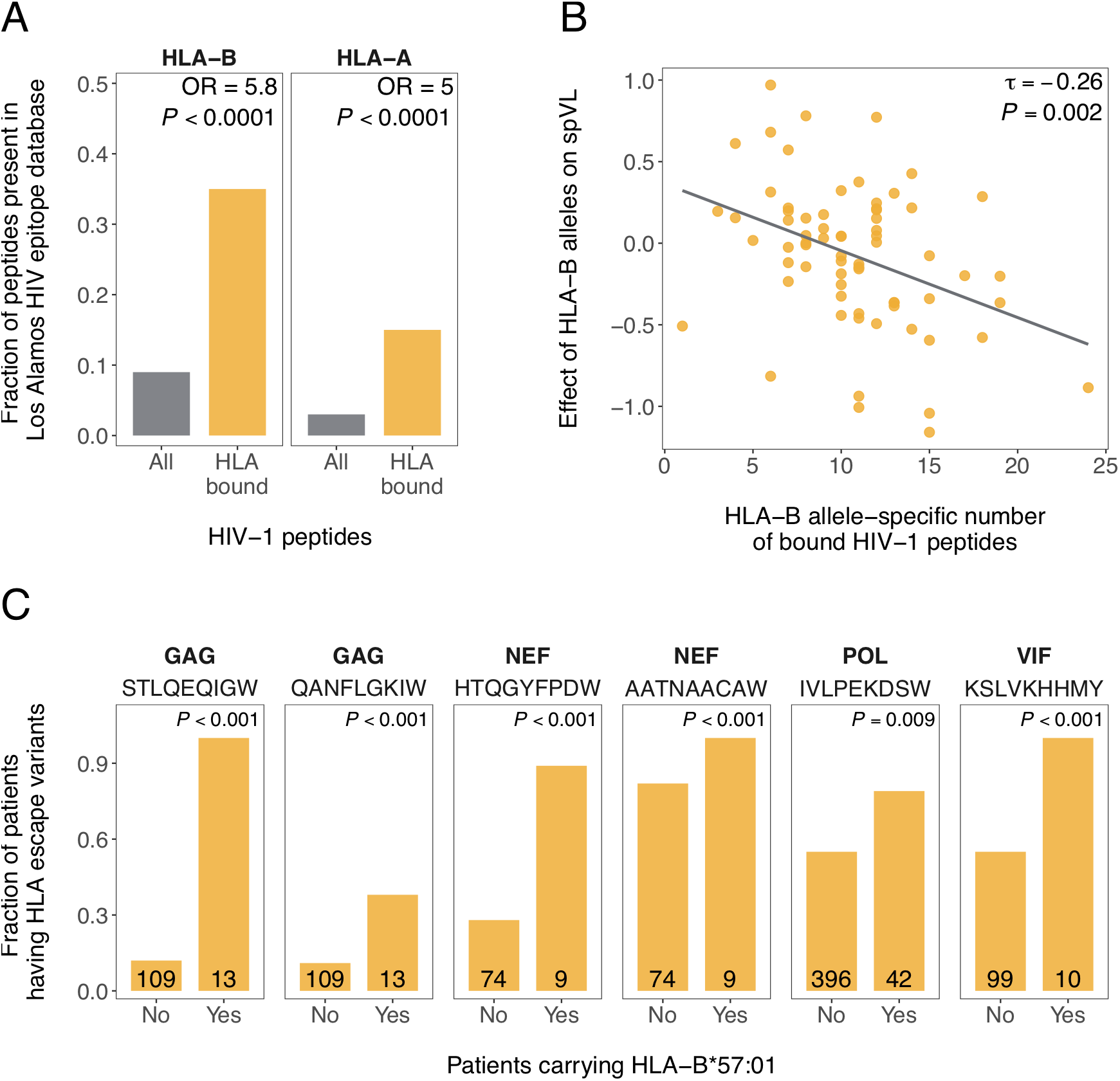
Characterization of predicted HLA-bound HIV-1 epitopes. (A) In comparison to all possible HIV-1 proteome-wide peptides (N = 3,252), the subsets of peptides predicted to be bound by HLA-B (N = 214) and HLA-A (N = 173), respectively, were strongly enriched for known epitopes defined in the Los Alamos HIV Molecular Immunology Database. Odds ratios (OR) and *P* values from Fisher exact test are shown. (B) HLA allele-specific effects on set point viral load (spVL) were calculated using linear regression and correlated with the predicted number of bound HIV-1 peptides. Each dot represents a distinct HLA-B allele (N = 69). Kendall correlation coefficient and p-value are shown. (C) For 6 of 18 reference peptides bound by HLA-B*57:01, the proportion of HLA escape variants in autologous sequence data differed significantly between B*57:01 carriers and non-carriers. The number of patients with available autologous HIV-1 sequence data is given inside each bar. Bonferroni-corrected *P* values from Fisher exact test are shown.

**Table 1.** Improvement in model fit by predicting epitopes from autologous HIV-1 sequences.

| HIV Protein | Number of Samples | % of non-missing sites in autologous sequences | Variation associated with epitopes from |  |
| --- | --- | --- | --- | --- |
|  |  |  | Reference sequence | Autologous sequence |
| *Gag* | 122 | 86.8 | 0 | 0.016 |
| *Pol* | 438 | 51.1 | 0.002 | 0.003 |
| *Vpu* | 103 | 88.7 | 0 | 0 |
| *Vpr* | 102 | 97.3 | 0 | 0 |
| *Nef* | 83 | 87.0 | 0.012 | 0.040 |
| *Gp41* | 83 | 86.0 | 0 | 0 |
| *Vif* | 110 | 94.5 | 0.050 | 0.051 |
| *Rev* | 109 | 84.1 | 0 | 0 |

In 4 out of 8 HIV-1 proteins for which autologous sequences were available, the variation (estimated as adjusted ΔR^2^) in set point viral load (spVL) associated with individual HLA-bound epitope repertoires increased when we predicted epitopes from autologous sequences instead of the HIV-1 reference sequence. Autologous sequence information was only available for a small subset of patients and coverage of the given protein sequences was incomplete. Estimates for both reference and autologous epitopes are therefore based on the same subset of patients and incomplete sequence data in order to make them comparable. No sequence data was available for the *Env* protein.

Next, we tested whether the patient-specific repertoire of predicted HIV-1 epitopes, defined by the number of peptides predicted to be bound by the specific HLA allele combination of the patient, was associated with spVL. For this, we ran a linear regression across the 6,315 HIV-1 patients. We first focused on known CTL epitopes from the Los Alamos HIV Molecular Immunology Database ^12^, of which 80 were represented among the 214 predicted HLA-B bound epitopes. The total number of these known CTL epitopes bound by patient-specific HLA-B variants accounted for only 1.8% variation in spVL (**Figure 2**). In contrast, the total number of all predicted HLA-B-bound epitopes per patient (including known and ‘uncharacterized’ epitopes) accounted for 5.3% variation in spVL (**Figure 2**). However, this was still lower than the 11.4% variation associated with genetic variation at HLA-B in previous genotype-based studies, suggesting that the total predicted epitope repertoire still included peptides irrelevant for the association between HLA and HIV-1. We thus aimed to refine the repertoire of predicted HLA-bound HIV-1 epitopes further to include only disease-relevant epitopes. For this, we calculated the epitope-specific association with spVL by running a separate linear regression for each predicted epitope and recording R^2^ and β-coefficient as measures of the epitope’s effect on spVL. This is analogous to the approach of a genome-wide association study (GWAS), where each genetic variant is tested for its association with a given trait, except that here we focus on functional protein variation (peptide binding by a patient’s HLA molecules) rather than genetic variation. Following this analogy, we term our approach *peptidome-wide association study* (PepWAS). Of 214 HIV-1 epitopes predicted to be bound by HLA-B, 132 accounted for nominal variation (adjusted R^2^ value > 0) in spVL, 74 of which were negatively and 58 positively associated with spVL (β-coefficients ranging from −0.1 to 0.77; **Table S2**). Importantly, we do not require statistical significance at this point as this is a candidate screen and we thus aim to minimize the number of false negatives. Subsequently, we designate the nominally associated epitopes as *disease-associated* predicted epitopes, even though their effects are not necessarily independent as they were tested with separate regression models. An analogous investigation of peptide binding by HLA-A alleles revealed an additional 74 disease-associated epitopes (**Table S3**).

**Figure 2.**
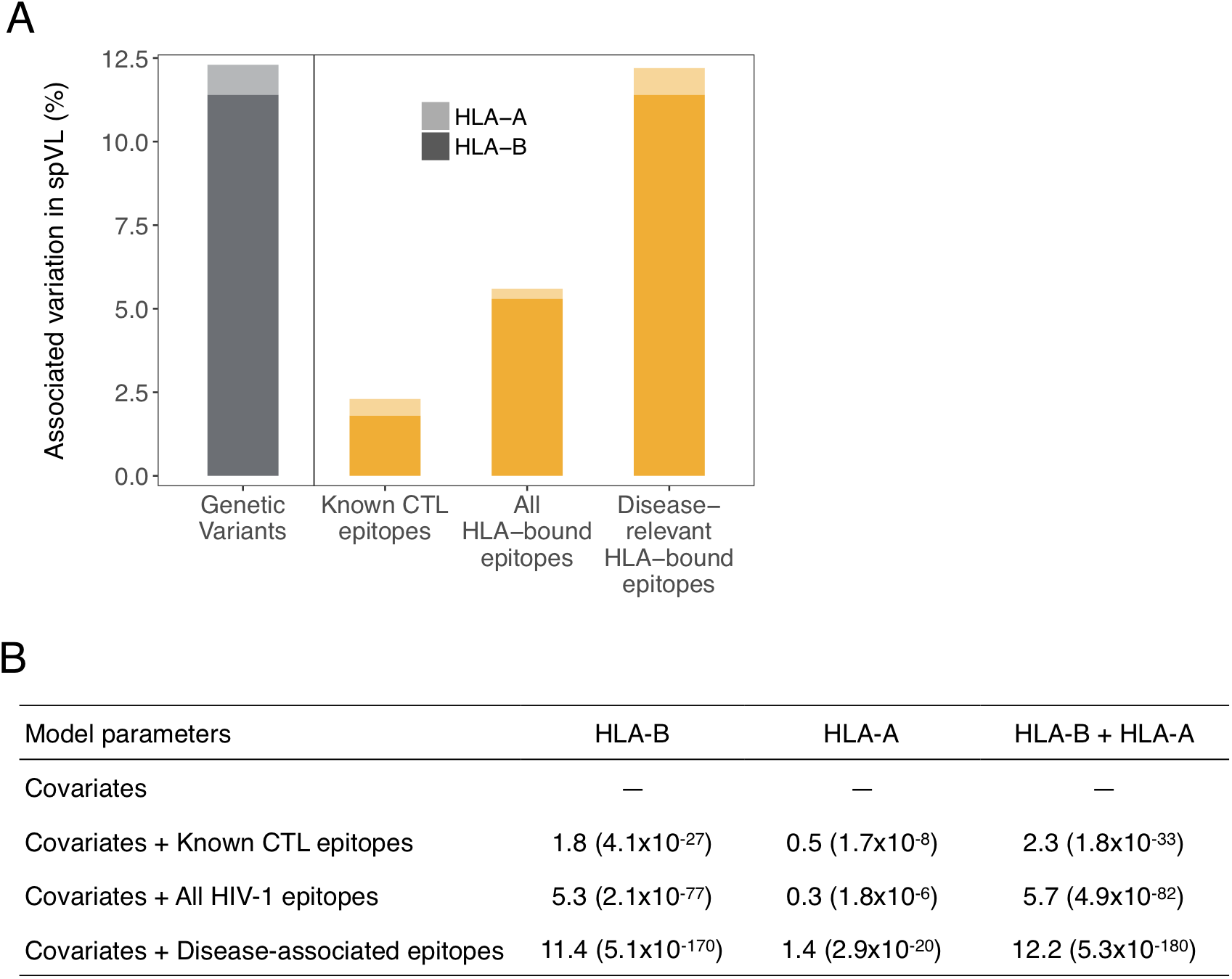
Variation in viral load associated with predicted epitope repertoires bound by HLA-B and HLA-A. Among HIV patients (N = 6,315), the proportion of variation (estimated as adjusted ΔR^2^) in set point viral load (spVL) associated with the patient-specific number of predicted HLA-bound HIV-1 epitopes is shown separately for HLA-B and HLA-A, and for different epitope sets. **(A)** Previously, 11.4% and 0.9% of the variation in spVL had been associated with independent genetic variants in HLA-B and HLA-A, respectively (grey bars; data from ref. 4). Here we instead calculated the variation in spVL associated with individual HLA-bound HIV epitope repertoires (yellow bars), based on known CTL epitopes from Los Alamos HIV Molecular Immunology Database, all HLA-bound HIV epitopes, and only the disease-associated HIV epitopes (the latter corresponding to 99.2% of the variation previously associated with genetic variation). **(B)** Variation associated with different sets of predicted epitopes. *P*-values (in parentheses) indicate the improvement over null model (covariates only: first five PCs and cohort group). Number of disease-associated predicted epitopes is 132 for HLA-B, and 74 for HLA-A, respectively.

Having refined the predicted HIV-1 epitope repertoire to only disease-associated epitopes, we then tested whether this subset accounted for a larger fraction of the variation in spVL than the total HIV-1 epitope repertoire. Indeed, the patients’ ability to bind a smaller or larger fraction of the HLA-B-specific disease-associated epitopes accounted for 11.4% of the variation in spVL (**Figure 2**). Similarly, the total number of predicted HIV-1 epitopes bound by individual HLA-A genotypes accounted for 0.3% of the variation, while disease-associated predicted epitopes accounted for 1.4% of the variation in spVL. On average, a patient’s HLA-B allele pair bound 16.2 ±7 (SD) disease-associated predicted HIV-1 epitopes, while its HLA-A alleles bound significantly less (6.6 ±6.5; Paired Wilcox rank sum test, *P* < 0.0001; **Figure S4**), potentially contributing to the stronger spVL-association of HLA-B compared to HLA-A. HLA-C-bound epitopes did not show any significant association with spVL, mirroring the lack of independent genetic associations for HLA-C in the latest GWAS ^4^. Predicted disease-associated epitopes of HLA-B and HLA-A together accounted for 12.2% of the variation in HIV-1 viral load, approximately corresponding to the 12.3% variation previously attributed to all independent genetic associations in the entire HLA (**Figure 2A**).

Interestingly, *Env* protein showed the largest number of disease-associated predicted epitopes, with both positive and negative effects. Among the disease-associated predicted HLA-B-bound epitopes, *Env*-derived epitopes alone accounted for 6.4% of variation in spVL, the highest among all HIV-1 proteins (**Figure 3A**). In addition to already known *Env*-derived epitopes associated with disease control e.g. RIKQIINMW, HRLRDLLLI ^22^, ERYLKDQQL ^23^, our analysis revealed previously undescribed HLA-epitope complexes e.g. B*57:01-STQLFNSTW, -NSTWFNSTW, or -RGWEALKYW showing strong negative associations with viral load (**Figure 3C**). Notably, several of the represented HLA alleles were predicted to bind both negatively and positively disease-associated epitopes (**Tables S4 and S5**), i.e. epitopes bound by the same HLA allele did not necessarily have the same effect on viral load.

**Figure 3.**
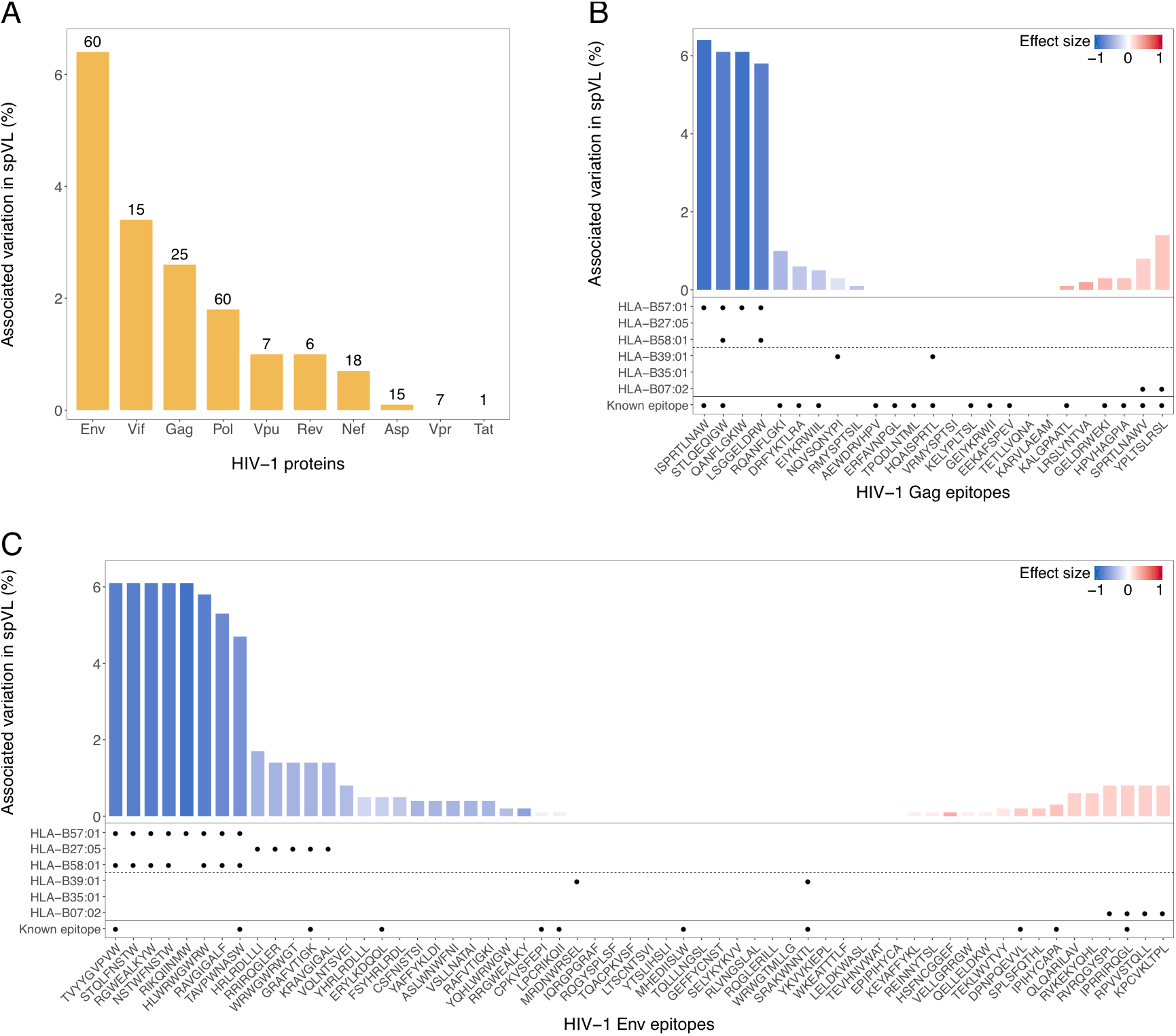
Epitope- and protein-specific association with viral load. **(A)** Percent of variation in spVL associated with all predicted epitopes of a given HIV-1 protein. Absolute number of predicted HLA-B bound epitopes per protein is shown above the bars. (B-C) Predicted HLA-B-bound epitopes accounted for varied levels of variation in set point viral load (spVL). Height of the bar represents the fraction of variation in spVL associated with each epitope, while the color reflects each epitope’s effect on spVL, ranging from protection (blue) to risk (red). Note that epitope effects are estimated separately and are thus not independent. *Gag* **(B)** and *Env* **(C)** proteins are shown as representative examples, together with information on predicted binding for 3 protective and 3 risk HLA-B alleles highlighted in a recent review ^35^ and whether peptides are known epitopes in Los Alamos HIV database. All other HIV-1 proteins are shown in Figure S3.

HLA molecule variants are known to bind peptide repertoires with distinct anchor motifs, based on the composition of their peptide-binding groove ^21^. This entailed the possibility that our PepWAS approach is merely identifying distinct groups of peptides per HLA variant, thus translating the known HLA variant-specific effect on viral load into peptide group-specific effects. While still helpful in guiding epitope research, this would provide only limited knowledge-gain compared to the HLA allele-specific associations known from previous work ^4^. In order to test for this possibility, we performed a cluster analysis on the predicted disease-associated epitopes bound by HLA-B (N = 132) and analyzed cluster-specific motifs and HLA allele binding patterns. Intriguingly, among the ten most dominant epitope clusters, each exhibiting a distinct peptide motif, nine were defined by multiple HLA-B alleles (**Figure 4**). All of these clusters included both novel and previously described epitopes, and three of them were defined by both risk- and protection-conferring alleles. Furthermore, all HLA variants bound peptides of multiple dominant clusters; e.g. B*57:01 is associated with 3 dominant clusters, each showing a distinct peptide motif, but all showing a strong preference for amino acid ‘W’ at anchor position 9 (**Figure 4**). Overall, the cluster analysis shows that our PepWAS approach identifies groups of peptides with distinct motifs that are different from HLA variant-specific binding motifs.

**Figure 4.**
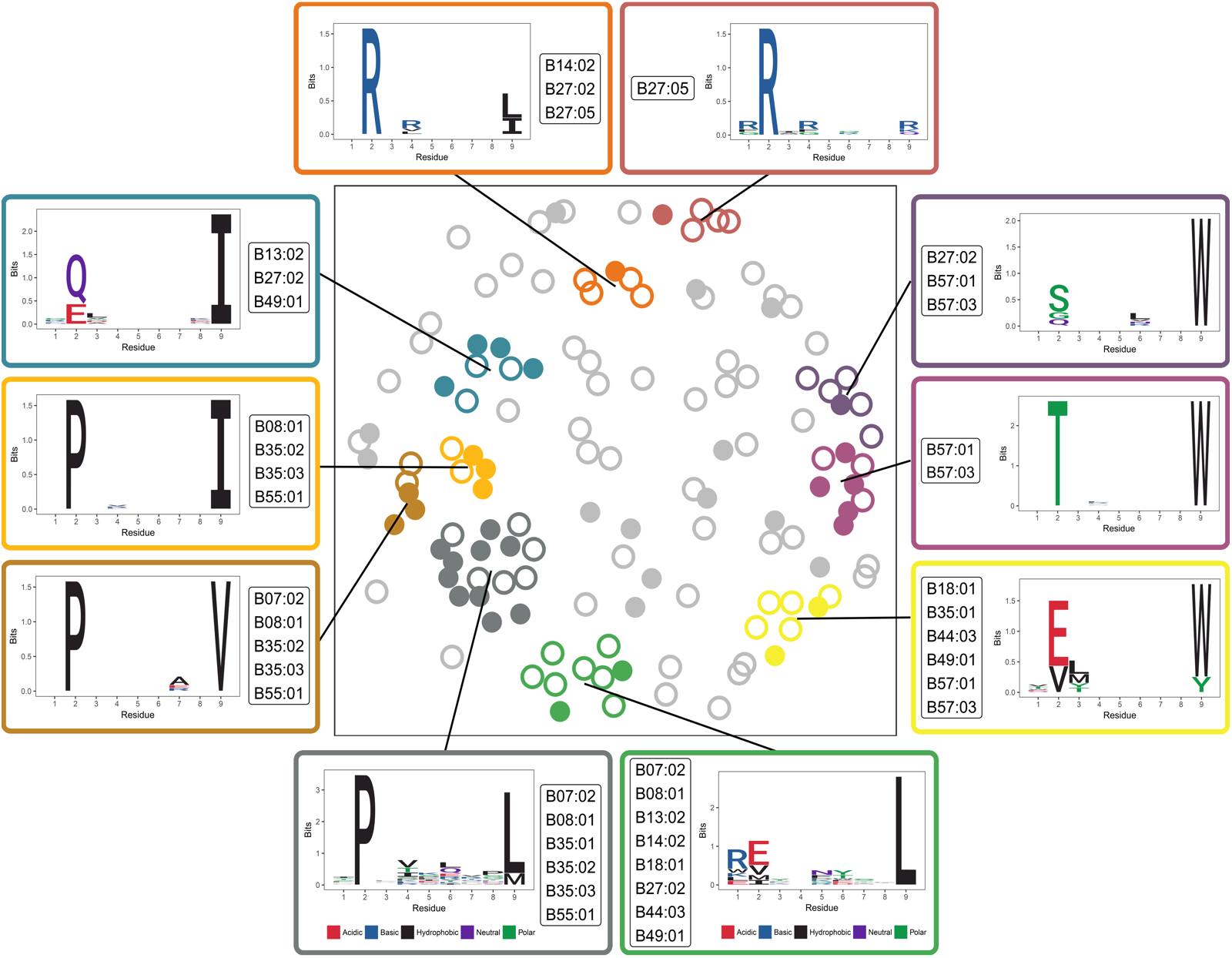
Clusters of disease-associated epitopes. Non-metric multidimensional scaling (NMDS) was used to visualize the pairwise distance between predicted HLA-B-bound disease-associated epitopes, which revealed 10 dominant clusters. Each circle represents an HLA-B bound disease-associated epitope (N = 132). Filled circles represent known CTL epitopes from the Los Alamos HIV Molecular Immunology Database (N = 45). Cluster-specific motif and HIV-1 associated HLA-B alleles (N = 16) binding the cluster’s epitopes are shown.

Generally, the 24 disease-associated epitopes predicted to be bound by HLA-B*57:01 (but also by other alleles), accounted for the highest level of variation in spVL, even though they derived from 5 different HIV-1 proteins (**Figure 3B, C and Figure S3**). One of these epitopes, the well characterized HIV-1 *Gag* epitope ISPRTLNAW (belonging to the violet cluster in **Figure 4**), slightly exceeded the effect of all other epitopes (**Figure 3B**), in concordance with experimental evidence ^24^. Predicted HIV-1 epitopes with a positive association to viral load, indicating disease risk, may represent peptide variants that fail to elicit a CTL response or which can readily mutate with negligible fitness cost for the virus. Indeed, the most risk-associated predicted *Vpu* epitope, IPIVAIVAL (**Figure S3F**; belonging to the largest, grey cluster in **Figure 4**), includes an anchor residue that exhibits significant variation in primary HIV-1 clones and is involved in mediating immune-evasion through down-regulation of HLA-C ^25^.

So far, our analysis was based on the HIV-1 genome reference sequence. Though widely used for research, focusing on this sequence accession may restrict our findings. We thus repeated the entire analysis using the HIV-1 proteome consensus sequence from the Los Alamos database, which incorporates major variation across different HIV-1 strains. The results remained qualitatively the same (**Figure S5 and Table S6**). However, HIV is well known to exhibit substantial within-host evolution ^26,27^ and it is easily conceivable that the ability of a patient’s HLA variants to bind HIV epitopes is significantly affected by genetic variation in the patient’s HIV population ^17^. We therefore also analyzed patient-specific autologous HIV-1 sequence information, which was available for a small subset of patients, covering 8 of the 10 HIV-1 proteins (**Table 1**). For 4 proteins (*Gag, Pol, Vif* and *Nef*) we found that HLA-bound epitope repertoires predicted from autologous sequences accounted for more variation in spVL than their homologs from the reference sequence (**Table 1**).

Overall, our findings reveal a functional basis of the robustly established association between HLA genes and HIV-1 infection outcome. We show that both quantity and quality of HLA-bound HIV epitopes contribute to controlling a patient’s viral load. Our data also suggests a more important role for *Env* protein-derived epitopes than previously thought. Ultimately, our PepWAS approach of combining computational HLA-specific epitope prediction with disease phenotype validation provides a promising avenue for identification and prioritization of novel epitopes as potential targets for immunotherapy. As such it may be applied to any HLA-associated complex disease.

## Materials and Methods

### Samples and Genotype data

We analyzed data of 6,311 chronically HIV-1 infected patients. The original data and thorough quality check are described in detail in McLaren *et al.* ^4^. Briefly, genome-wide genotype data had been collected from 8 independent GWAS studies and combined as part of the International Collaboration for the Genomics of HIV. Principal component analysis was used to infer ancestry using HapMap 3 ^28^ as a reference. Only samples grouping with HapMap Europeans were retained to ensure consistent ancestry across samples. The first five principal components were used as covariates to account for residual population stratification in downstream analysis. SNP genotypes that had not been covered in original genotyping platforms were imputed with Minimac ^29^ using haplotypes from the 1,000 Genomes Project Phase 1 v3 reference panel. Imputed genotypes from other imputation protocols, Shapeit ^30^ and Impute2 ^31^, yielded highly concordant results. Imputed SNPs with low r2 score (< 0.3) or minor allele frequency of < 0.5% were discarded. HLA alleles (best-guess genotypes at 4-digit resolution) for classical class I loci (HLA-A, -B, -C) were imputed from genome-wide genotype data using SNP2HLA and a reference panel of 5,225 individuals of European ancestry ^32^. Overall, 69 and 37 classical alleles for HLA-B and HLA-A, respectively, were represented in the data (Figure S1). Available measurement of pre-treatment set point viral load (spVL; log10 HIV-1 RNA copies/μl of plasma) was used as quantitative disease phenotype for all patients ^4^. All participants were HIV-1-infected adults, and written informed consent for genetic testing was obtained from all individuals as part of the original study in which they were enrolled ^4^.

### HLA binding affinity for HIV-1 epitopes

We used the NCBI accession NC_001802.1 as the reference sequence for the HIV-1 proteome (M group subtype B). It comprised 10 proteins with sequence length ranging from 82 to 1435 amino acids (Table S7). For the consensus sequence-based analysis, the most recent Consensus HIV-1 proteome (subtype B, year 2004, ID 104CP2) was taken from Los Alamos HIV sequence database. The *Gag-Pol* protein is a precursor protein that results from a −1 ribosomal frameshifting event in upstream *Gag* ^33^. It is cleaved by virus-encoded protease to produce the mature Pol protein. In our analysis, we manually trimmed the *Gag-Pol* protein sequence to Pol in order to avoid redundancy with the separate *Gag* protein. HLA class-I molecules preferentially bind and present 9mer peptides ^21,34^. NetMHCcons-1.1 is a computational method that predicts binding affinity of 9mer peptides to any known HLA class I molecule ^6^. It is a consensus approach which integrates three prediction methods, namely NetMHC 3.4, NetMHCpan 2.8 and PickPocket 1.1. The analyzed dataset includes some HLA alleles with no available training data for the prediction methods, which makes the consensus approach more favorable over either of the underlying methods ^6^. It also reports the rank of predicted binding affinity of HLA-peptide complexes against predicted affinity of 200,000 random natural peptides. Using this tool, we predicted HLA allele-specific binding affinities for all 9mer peptides generated from the entire HIV-1 proteome. HLA-peptide complexes with predicted binding affinity rank less than 0.5 were retained (corresponding to ‘strongly bound’ peptides ^6^). The breadth of peptides bound by a patient’s HLA allele pair was taken as the total number of unique peptides predicted to be bound by both alleles but accounting for multiple occurrences of peptides in the HIV-1 proteome. The specificity of the correlation between HLA-B allele-specific effect on spVL and HLA-B-bound HIV-1 peptides was confirmed by running the same correlation on predicted HLA-B-bound peptides from four randomly selected viruses, namely Dengue (NC_002640.1), Rhinovirus-A (NC_001617.1), Hepatitis-B (NC_003977.2) and Rubella (NC_001545.2).

### Association with viral load

The association of an allele or a peptide with viral load (spVL) was calculated using a linear regression model corrected for population covariates following McLaren *et al.* ^4^. Covariates included the first five principle components of SNP variation and the cohort identity (all adopted from McLaren *et al.* ^4^). Variation in viral load attributable to a given variable (allele or peptide) was calculated as the difference between adjusted-R2 values of the model with variable and covariates (Equation 1) and the model with covariates only (Equation 2), following McLaren et al. ^4^. The variable’s regression coefficient (β1) was used as the measure of its effect on viral load. ε, residual term, is the difference between predicted and true values of spVL given the value of variable and covariates.

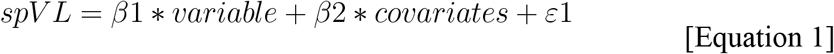

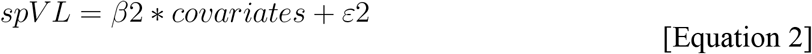

We selected 6 HLA-B alleles, highlighted by McLaren & Carrington ^35^, for representation in peptide-specific association plots (Figures 3B, C, S3 and S5). Three of them, namely B*57:01, B*58:01, and B*27:05, are associated with lower spVL, while the 3 others, namely B*07:02, B*35:01, and B*39:01 are associated with increased spVL. All analyses were performed in R v3.3.3 and data was visualized using the ggplot2 v2.2.1 package ^36^.

### Clustering of HLA-B bound epitopes

Position-associated entropy was calculated for all HLA-B-bound disease-associated epitopes (N = 132) using HDMD v1.2 package ^37^. Subsequently, entropy-weighted pairwise distance between those epitopes was calculated by using PMBEC similarity matrix as defined in Kim *et al.* ^38^. Non-metric multidimensional scaling was performed on a pairwise distance matrix using the ecodist v2.0.1 package ^39^. Density-based clustering was performed using the dbscan v1.1-1 package ^40^. For each cluster, HLA-B alleles with an overall significant association to viral load (*P*-value <= 0.05/69; corrected for multiple testing) are shown (N = 16). Sequence logo plots to represent cluster motifs were made using the ggseqlogo v0.1 package ^41^.

### HLA binding of peptides from autologous HIV-1 sequences

We analyzed autologous HIV-1 sequences from Bartha et al. ^16^. Autologous sequences were available for 8 of 10 HIV-1 proteins (only Gp41 segment for *Env*) but only for a small subset of patients in our cohorts (Table 1). Due to the bulk sequencing of viral RNA from pre-treatment stored plasma with limited coverage, there were large segments of missing sites in individual sequences, further reducing the available sequence information for epitope prediction. For comparison of autologous and reference epitopes, for each individual we extracted the corresponding parts of the reference sequence for which autologous sequence data was available. HLA-B binding affinities were calculated for autologous sequences and homologous reference parts using NetMHCcons-1.1 ^6^. HLA-peptide complexes with binding affinity rank <0.5 were retained (same criterion as used for the general analysis). The stronger association of autologous epitopes with viral load, compared to reference-based epitopes, further substantiates our epitope prediction pipeline and corroborates that within-host variation of HIV-1 can significantly alter the efficiency of HLA-based immunity.

### HLA-B allele specific escape mutations in HIV-1 epitopes

Of a total of 24 epitopes from the HIV-1 reference sequence that were predicted to be bound by HLA-B*57:01, autologous sequence information was only available for 18. For each of these 18 epitopes, we created a contingency table with the number of patients who are carriers and non-carriers of the HLA-B*57:01 allele, and the number of patients in whom the autologous version of the given epitope is and is not bound by B*57:01. Fischer exact test was performed on the contingency table to calculate the significance.

## Acknowledgements

Patient and HIV sequence data was collected and generously provided by the International Collaboration for the Genomics of HIV. This project has been funded in whole or in part with federal funds from the Frederick National Laboratory for Cancer Research, under Contract No. HHSN261200800001E. The content of this publication does not necessarily reflect the views or policies of the Department of Health and Human Services, nor does mention of trade names, commercial products, or organizations imply endorsement by the U.S. Government. This Research was supported in part by the Intramural Research Program of the NIH, Frederick National Lab, Center for Cancer Research as well as the Emmy Noether Programme of the Deutsche Forschungsgemeinschaft (grant LE 2593/3-1 to T.L.L.).

## Author Contributions

T.L.L. conceived the study. J.A., T.L.L., P.J.M. and J.F. designed the analyses. P.J.M., J.F. and N.C. provided data. J.A. and T.L.L. analyzed the data. J.A., T.L.L., P.J.M., J.F. and M.C. interpreted the results. J.A. and T.L.L. drafted the manuscript with critical feedback from M.C., P.J.M. and J.F.

## References

1. Goulder, P.J.R., and Walker, B.D. (2012). HIV and HLA Class I: An Evolving Relationship. Immunity 37, 426–440.

2. Fellay, J., Shianna K. V, Ge, D., Colombo, S., Ledergerber, B., Weale, M., Zhang, K., Gumbs, C., Castagna, A., Cossarizza, A., et al. (2007). A whole-genome association study of major determinants for host control of HIV-1. Science (80-.). 317, 944–947.

3. The International HIV Controllers Study, Pereyra, F., Jia, X., McLaren, P.J., Telenti, A., de Bakker, P.I.W., Walker, B.D., Team, A., Jia, X., McLaren, P.J., et al. (2010). The Major Genetic Determinants of HIV-1 Control Affect HLA Class I Peptide Presentation. Science 330, 1551–1557.

4. McLaren, P.J., Coulonges, C., Bartha, I., Lenz, T.L., Deutsch, A.J., Bashirova, A., Buchbinder, S., Carrington, M.N., Cossarizza, A., Dalmau, J., et al. (2015). Polymorphisms of large effect explain the majority of the host genetic contribution to variation of HIV-1 virus load. Proc. Natl. Acad. Sci.

5. Mellors, J.W., Rinaldo, C.R., Gupta, P., White, R.M., Todd, J. a, and Kingsley, L. a (1996). Prognosis in HIV-1 infection predicted by the quantity of virus in plasma. Science 272, 1167–1170.

6. Karosiene, E., Lundegaard, C., Lund, O., and Nielsen, M. (2012). NetMHCcons: A consensus method for the major histocompatibility complex class i predictions. Immunogenetics 64, 177–186.

7. Zhang, H., Lundegaard, C., and Nielsen, M. (2009). Pan-specific MHC class I predictors: A benchmark of HLA class I pan-specific prediction methods. Bioinformatics 25, 83–89.

8. Rooney, M.S., Shukla, S.A., Wu, C.J., Getz, G., and Hacohen, N. (2015). Molecular and genetic properties of tumors associated with local immune cytolytic activity. Cell 160, 48–61.

9. Strønen, E., Toebes, M., Kelderman, S., van Buuren, M.M., Yang, W., van Rooij, N., Donia, M., Böschen, M.-L., Lund-Johansen, F., Olweus, J., et al. (2016). Targeting of cancer neoantigens with donor-derived T cell receptor repertoires. Science (80-.). 352, 1337–1341.

10. Košmrlj, A., Read, E.L., Qi, Y., Allen, T.M., Altfeld, M., Deeks, S.G., Pereyra, F., Carrington, M., Walker, B.D., and Chakraborty, A.K. (2010). Effects of thymic selection of the T-cell repertoire on HLA class I-associated control of HIV infection. Nature 465, 350–354.

11. Martoglio, B., Graf, R., and Dobberstein, B. (1997). Signal peptide fragments of preprolactin and HIV-1 p-gp160 interact with calmodulin. EMBO J. 16, 6636–6645.

12. Yusim, K., Korber, B.T.M., Brander, C., Haynes, B.F., Koup, R., Moore, J.P., Walker, B.D., and Watkins, D.I. (2009). HIV molecular immunology. Los Alamos, New Mex. Los Alamos Natl. Lab. Theor. Biol. Biophys. 3–24.

13. Addo, M.M., Yu, X.G., Rathod, A., Cohen, D., Eldridge, R.L., Strick, D., Johnston, M.N., Corcoran, C., Wurcel, A.G., Fitzpatrick, C.A., et al. (2003). Comprehensive epitope analysis of human immunodeficiency virus type 1 (HIV-1)-specific T-cell responses directed against the entire expressed HIV-1 genome demonstrate broadly directed responses, but no correlation to viral load. J. Virol. 77, 2081–2092.

14. Goulder, P.J., Bunce, M., Krausa, P., McIntyre, K., Crowley, S., Morgan, B., Edwards, A., Giangrande, P., Phillips, R.E., and McMichael, a J. (1996). Novel, cross-restricted, conserved, and immunodominant cytotoxic T lymphocyte epitopes in slow progressors in HIV type 1 infection. AIDS Res. Hum. Retroviruses 12, 1691–1698.

15. Kiepiela, P., Ngumbela, K., Thobakgale, C., Ramduth, D., Honeyborne, I., Moodley, E., Reddy, S., de Pierres, C., Mncube, Z., Mkhwanazi, N., et al. (2007). CD8+ T-cell responses to different HIV proteins have discordant associations with viral load. Nat. Med. 13, 46–53.

16. Bartha, I., Carlson, J.M., Brumme, C.J., McLaren, P.J., Brumme, Z.L., John, M., Haas, D.W., Martinez-Picado, J., Dalmau, J., López-Galíndez, C., et al. (2013). A genome-to-genome analysis of associations between human genetic variation, HIV-1 sequence diversity, and viral control. Elife 2, 1–16.

17. Kawashima, Y., Pfafferott, K., Frater, J., Matthews, P., Payne, R., Addo, M., Gatanaga, H., Fujiwara, M., Hachiya, A., Koizumi, H., et al. (2009). Adaptation of HIV-1 to human leukocyte antigen class I. Nature 458, 641–645.

18. Phillips, R.E., Rowland-Jones, S., Nixon, D.F., Gotch, F.M., Edwards, J.P., Ogunlesi, A.O., Elvin, J.G., Rothbard, J.A., Bangham, C.R.M., Rizza, C.R., et al. (1991). Human immunodeficiency virus genetic variation that can escape cytotoxic T cell recognition. Nature 354, 453–459.

19. Roberts, H.E., Hurst, J., Robinson, N., Brown, H., Flanagan, P., Vass, L., Fidler, S., Weber, J., Babiker, A., Phillips, R.E., et al. (2015). Structured observations reveal slow HIV-1 CTL escape. PLoS Genet. 11, e1004914.

20. Kaslow, R.A., Carrington, M., Apple, R., Park, L., Munoz, A., Saah, A.J., Goedert, J.J., Winkler, C., O’Brien, S.J., Rinaldo, C., et al. (1996). Influence of combinations of human major histocompatibility complex genes on the course of HIV-1 infection. Nat Med 2, 405–411.

21. Falk, K., Rötzschke, O., Stevanovié, S., Jung, G., and Rammensee, H.-G. (1991). Allele-specific motifs revealed by sequencing of self-peptides eluted from MHC molecules. Nature 351, 290–296.

22. Pereyra, F., Heckerman, D., Carlson, J.M., Kadie, C., Soghoian, D.Z., Karel, D., Goldenthal, A., Davis, O.B., DeZiel, C.E., Lin, T., et al. (2014). HIV control is mediated in part by CD8+ T-cell targeting of specific epitopes. J. Virol. 88, 12937–12948.

23. Borrow, P., Lewicki, H., Hahn, B.H., Shaw, G.M., and Oldstone, M.B.A. (1994). Virus-Specific CD8+ Cytotoxic T-Lymphocyte Activity Associated with Control of Viremia in Primary Human Immunodeficiency. 68, 6103–6110.

24. Llano, A., Williams, A., Olvera, A., Silva-Arrieta, S., and Brander, C. (2013). Best-Characterized HIV-1 CTL Epitopes: The 2013 Update. HIV Mol. Immunol. 2013 3–25.

25. Apps, R., Del Prete, G.Q., Chatterjee, P., Lara, A., Brumme, Z.L., Brockman, M.A., Neil, S., Pickering, S., Schneider, D.K., Piechocka-Trocha, A., et al. (2016). HIV-1 Vpu Mediates HLA-C Downregulation. Cell Host Microbe 19, 686–695.

26. Cotton, L.A., Kuang, X.T., Le, A.Q., Carlson, J.M., Chan, B., Chopera, D.R., Brumme, C.J., Markle, T.J., Martin, E., Shahid, A., et al. (2014). Genotypic and Functional Impact of HIV-1 Adaptation to Its Host Population during the North American Epidemic. PLoS Genet. 10,.

27. Li, G., Piampongsant, S., Faria, N.R., Voet, A., Pineda-Peña, A.-C., Khouri, R., Lemey, P., Vandamme, A.-M., and Theys, K. (2015). An integrated map of HIV genome-wide variation from a population perspective. Retrovirology 12, 18.

28. The International HapMap 3 Consortium (2010). Integrating common and rare genetic variation in diverse human populations. Nature 467, 52–58.

29. Howie, B., Fuchsberger, C., Stephens, M., Marchini, J., and Abecasis, G.R. (2012). Fast and accurate genotype imputation in genome-wide association studies through prephasing. Nat. Genet. 44, 955–959.

30. Delaneau, O., Zagury, J.-F., and Marchini, J. (2013). Improved whole-chromosome phasing for disease and population genetic studies. Nat. Methods 10, 5–6.

31. Howie, B.N., Donnelly, P., and Marchini, J. (2009). A flexible and accurate genotype imputation method for the next generation of genome-wide association studies. PLoS Genet 5, e1000529.

32. Jia, X., Han, B., Onengut-Gumuscu, S., Chen, W.M., Concannon, P.J., Rich, S.S., Raychaudhuri, S., and de Bakker, P.I.W. (2013). Imputing Amino Acid Polymorphisms in Human Leukocyte Antigens. PLoS One 8,.

33. Jacks, T., Power, M.D., Masiarz, F.R., Luciw, P.A., Barr, P.J., and Varmus, H.E. (1988). Characterization of ribosomal frameshifting in HIV-1 gag-pol expression. Nature 331, 280–283.

34. York, I.A., Chang, S.-C., Saric, T., Keys, J.A., Favreau, J.M., Goldberg, A.L., and Rock, K.L. (2002). The ER aminopeptidase ERAP1 enhances or limits antigen presentation by trimming epitopes to 8-9 residues. Nat. Immunol. 3, 1177–1184.

35. McLaren, P.J., and Carrington, M. (2015). The impact of host genetic variation on infection with HIV-1. Nat. Immunol. 16, 577–583.

36. Wickham, H., and Wickham, M.H. (2007). The ggplot package.

37. McFerrin, L., and McFerrin, M.L. (2013). Package HDMD. Stazeno Z Http://cran.R-Project.org/web/packages/HDMD/HDMD.Pdf (14.6. 2013).

38. Kim, Y., Sidney, J., Pinilla, C., Sette, A., and Peters, B. (2009). Derivation of an amino acid similarity matrix for peptide:MHC binding and its application as a Bayesian prior. BMC Bioinformatics 10, 394.

39. Goslee, S., and Urban, D. (2007). The ecodist Package: Dissimilarity-based functions for ecological analysis. J Stat Softw 22, 1–19.

40. Hahsler, M., and Piekenbrock, M. (2017). dbscan: Density Based Clustering of Applications with Noise (DBSCAN) and Related Algorithms. R Packag. Version 0-1.

41. Wagih O. (2017). ggseqlogo: a versatile R package for drawing sequence logos. Bioinformatics 33, 3645–3647.

